# Winners and losers under past and future climate change

**DOI:** 10.1101/2023.09.11.556540

**Authors:** Anne E. Thomas, Matthew J. Larcombe, Steven I. Higgins, Antonio Trabucco, Andrew J. Tanentzap

## Abstract

Understanding the historical and physiological context of species’ vulnerabilities to climate change is a crucial step in predicting “winners” and “losers” under climate change. However, few studies have compared the magnitude and mechanisms of extant species’ responses to climate change in both the past and the future. By combining temporally contrasting range and niche projections, we show that range shifts in the next 50 years will need to be more extreme than in the past 6000 years to track climate niches in a large plant radiation. A new subset of physiological niche traits, particularly temperature and radiation tolerance, will be strong filters of range occupancy under anthropogenic compared with Holocene climate change. In the absence of migration, temperature niche shifts tracking the magnitude of climate change will also be required for many species to maintain their present ranges. Where range shifts occur, our results suggest that communities will be restructured differently in different habitats, with widespread range contraction in the mountains and potential latitudinal range expansion in the lowlands. Our study adds to a growing body of evidence that despite the threats posed by climate change to many species, not all species will experience unmitigated loss, and that it may be possible to predict which species are most at risk based on physiological and geographical traits.

## INTRODUCTION

Climate change is putting unprecedented pressure on plants to adapt or migrate to avoid extinction. Global temperatures are already warmer than the last sustained warm period 6500 years ago and are projected to increase from preindustrial levels by up to 5.7 °C by 2100, the highest global temperature in millions of years (Foster et al., 2017; IPCC, 2021). Evidence of climate-driven range shifts has been found in the past (e.g., Davis & Shaw, 2001) and present (e.g., Chen et al., 2011). Presently, these shifts are largest in areas with the greatest warming (Chen et al., 2011). *In situ* adaptation is also occurring at the pace of current climate change (Bradshaw & Holzapfel, 2006; Davis et al., 2005). Adaptation is arguably unlikely to preclude the need for migration because niche conservatism is widespread (Wiens et al., 2010) and there are other barriers to rapid adaptive evolution (Mokany et al 2019, Kremer et al 2012, Visser 2008, Franks 2013, Merila and Hendry 2013). Quantifying the vulnerability of species to future climate change via their potential to migrate or adapt remains an unresolved challenge in biodiversity conservation, as conservation efforts hinge on whether measures such as protected areas or assisted migration will be adequate for a given taxa.

Species vary in their vulnerability to climate change (Nadeau et al., 2017; Thomas et al., 2011). Some species may experience more pressure to move or adapt due to the narrowness of their environmental tolerances, or habitat-and region-specific exposure to climate change (Antão et al., 2022; Nogués-Bravo et al., 2018; Williams et al., 2008). Other species may experience little pressure due to broad tolerances or plasticity (Corlett & Westcott, 2013), and some may even experience net expansion of potential range, thus potentially benefiting from climate change (Thomas et al., 2011). An important goal of predicting the future vulnerability to climate change is identifying what characteristics distinguish potential “winners” and “losers” under climate change (Araújo et al., 2011). For example, Thuiller et al. (2005) found that projected range stability under future climate change could be predicted by niche characteristics such as breadth and marginality. Connecting vulnerability to physiological niche traits mechanistically linked to plant growth would take predicting climate change impacts a step beyond correlative environmental axes.

One limitation of measuring the vulnerability of species only to future climate change is that it overlooks background changes happening over longer timescales (Nogués-Bravo et al., 2018). Extant species have all experienced climate fluctuation in the recent geological past (Pardi & Smith, 2012; Wanner et al., 2008), which can influence the magnitude and mechanisms of their responses to future change (Nadeau et al., 2017). For example, Liang et al. (2018) showed that alpine plant ranges in the Hengduan Mountains have shifted farther since the last glacial maximum than they are projected to shift by 2050, but that future shifts will occur at greater velocity. On an evolutionary scale, Jezkova & Wiens (2016) found that to keep pace with climate change, niche shifts along temperature axes must be orders of magnitude higher than the rate of past niche change relative to species’ most recent common ancestors. However, few studies have compared both range and niche shifts of extant species in the past and future. Alongside understanding differences in the magnitude of changes, such comparisons can help determine whether vulnerability to change can be consistently predicted by niche traits across time. If so, species’ past niche responses may be able to help identify future climate change winners and losers and inform appropriate conservation action. If not, the contrast between past and future niche responses can focus conservation efforts on species requiring the most extreme responses.

The goal of this study was to compare range and niche sensitivity between past and future climate to identify predictors of plant responses to change useful for conservation. To do so, we quantified niche and range change in a large plant radiation endemic to New Zealand, *Veronica* sect. *Hebe.* The radiation consists of 124 species that occupy a wide variety of habitats and range sizes (Meudt et al., 2015). The size and shared history of the clade makes it an ideal case study for tracing the effects of climate change on range and niche stability across species with different environmental tolerances. We estimated species’ environmental tolerances to Holocene and future climate change using a physiological model of plant growth and resource use (Higgins et al., 2012). The model allows both niche traits and potential range change to be estimated under past, present-day, and future climates. With this approach, we asked: 1) How do predicted range shifts compare between periods of past and future climate change? 2) Which niche traits best predict range change at different time periods? 3) How much would niche traits need to shift for species’ current ranges to match environmental conditions in each time period? Answering these questions provides temporal and physiological context for efforts to predict climate change impacts on plant species.

## METHODS

### Physiological niche modeling

We quantified species’ environmental niches with 24 physiological traits estimated by a process-based species distribution model (SDM) derived from the Thornley Transport Resistance (TTR) model of plant growth (Higgins et al., 2012). The TTR SDM estimates carbon and nutrient uptake and allocation given temperature, solar radiation, soil moisture, and soil nitrogen data associated with species occurrences. The resulting niche traits for each species are defined by upper and lower environmental limits on physiological processes, such as the temperature that constrains photosynthesis, soil moisture concentration that constrains nitrogen uptake, and temperature that constrains growth (see Table S1 for details). These niche traits can be used to project the suitable ranges of the species, and the parameters defining them also represent niche traits in downstream analyses (e.g. Larcombe et al., 2018). Because these modeled TTR traits can range beyond realistic physiological limits where the species’ range is more sensitive to other co-limiting variables (Higgins et al., 2012), we additionally set minimum and maximum boundaries on temperatures-based traits. For the temperature limiting photosynthesis (represented in TTR by maximum monthly temperature), we set the lower bound at -5 °C and the upper bound at 40 °C (Kramer & Kozlowski, 1979). For the temperature limiting growth (represented by monthly minimum temperature), the bounds were set at 0 °C and 25 °C (Lambers et al., 1998). For the temperature limiting nitrogen uptake and respiration (represented by monthly mean temperature), the bounds were set at 0 °C and 30 °C (Lambers et al., 1998). Within these limits, some traits will still not reflect a precise response curve for the target process when a species’ range is more constrained by other variables. Instead, they reflect the sensitivity of the projected range to the target process at a given temperature and so can still enable meaningful relative comparisons between species. Nevertheless, to avoid diluting the effect of more sensitive parameters, we removed 7 traits from the downstream analysis for which more than half of the species had values set at the trait boundaries, leaving 17 traits.

We fitted the SDM to each of 84 species using geo-referenced herbarium records curated for an e-flora treatment of *Veronica* by Phil Garnock-Jones. A total of 9167 georeferenced *Veronica* occurrences were used, with 19 to 586 records per species (mean: 98.6, median: 66; Table S2). Environmental data for modern climate included temperature estimates from WorldClim (1960-1990 mean; Hijmans et al., 2005), soil nitrogen (Shangguan et al., 2014), and solar radiation and soil moisture from the CGIAR database (Trabucco & Zomer, 2010), all at 1-km resolution. We generated random pseudoabsence points equal in number to occurrences after Larcombe et al. (2018). Pseudoabsence points were distributed proportionally across ten environmental zones classified from the environmental input variables with the CLARA algorithm in the R package cluster v2.07. Although the TTR SDM models biomass-based abundance values, abundance is affected by factors not included in the model, such as herbivory. To avoid unrealistic assumptions, abundance values are transformed into probabilities of presence and absence (Higgins et al., 2012). Binary presence-absence is then predicted based on the probability threshold that maximizes the proportions of true positive and true negative predictions. To allow model evaluation, 25% of the presence and pseudoabsence points were withheld as testing data. Evaluation metrics included false positive and false negative rates and the area under the receiver operating characteristic curve (AUC).

We then estimated how projected species ranges changed under past and future climate scenarios in the absence of niche shifts. Using the TTR niches estimated from the current climate, we projected suitability in New Zealand for each species in the mid-Holocene climate (“paleo”, about 6000 years ago) and in 2070 under two future climate change scenarios. Future climate scenarios included the RCP 4.5 (intermediate mitigation) and RCP 8.5 (worst case) CO2 concentration pathways. Paleo and future temperature projections were downscaled and calibrated by WorldClim from the Coupled Model Intercomparison Project 5 data at 1-km resolution. Soil moisture data were derived from these paleo and future climate scenarios after Trabucco and Zomer 2010 using a model of evapotranspiration. To calculate range changes between time periods, we calculated the overlap of grid cells where species were predicted to be present between both the projected paleo and present ranges, and between the present and each future scenario range. Non-overlapping cells were categorized as lost or gained range, and percent of original range lost or gained was calculated for each of the comparisons: paleo to current, current to RCP 4.5, current to RCP 8.5 (each subsequently referred to by the focal time period).

To compare potential species niche adaptation to past and future climate in the absence of migration, we fitted the TTR SDM independently using mid-Holocene, RCP 4.5, and RCP 8.5 climate scenarios, following the same protocol as above. Given that species have possibly migrated to track climate niche in the last 6000 years (Corlett & Westcott, 2013; Davis & Shaw, 2001; Pardi & Smith, 2012), these models are not meant to represent accurate paleo-niches <u>(see Nogués-Bravo, 2009)</u>, but are a conceptual reference point for how much niche change would be required to maintain the same present-day range. Within the next 50 years, lags in migration may limit movement (Bertrand et al., 2011), but the same logic applies.

### Statistical analysis

To compare range loss and range gain between time periods and identify which traits were most strongly associated with range change, we fitted phylogenetic linear mixed models with the R package phyr v1.1.0. We first reduced the niche space and eliminated correlated traits by selecting a subset of TTR niche traits based on a phylogenetic PCA. We determined which TTR parameters explained the most variation in the clade by examining the first 7 principal component axes, which collectively explained 95% of the variance (Table S3). We selected traits with the highest eigenvector loadings for each principal component based on the threshold of a single variable’s expected proportion of the sum of squares (Table S3). Finally, biplots were examined visually to remove correlated traits. This reduced the trait space to 9 parameters (Table S1, S3). We then mean-centered and scaled TTR traits by their standard deviations. The models were fitted using Bayesian Markov chain Monte Carlo simulations with default, weakly informative INLA priors. The percent of original range lost or gained was modelled as a Gaussian response, and predictors included the nine selected TTR traits, habitat (mountain, lowland, or both), and interactions of each trait with time period (paleo, RCP4.5, and RPC8.5), along with the main effects of each period. Habitat was determined with elevation data from herbarium records, field guides and floras (Bayly & Kellow, 2006; NZPCN, 2021), and expert advice (P. Garnock-Jones, pers. comm.). Generally, mountain habitat corresponded to >400 m a.s.l., varying with latitude (Wardle, 1991). Random effects included a non-phylogenetic species effect, accounting for repeated measurements in each time period, and the phylogenetic covariance of species. Range gain was quarter-root transformed to reduce the influence of outliers. Effects were considered statistically significant when the 95% confidence interval of the posterior distribution excluded 0. We also separately tested for phylogenetic signal in range loss and gain by calculating Blomberg’s K and Pagel’s lambda with the R package phytools v0.7. All analyses were performed with the phylogenetic tree estimated in Chapter 3, pruned to the species that had enough occurrences for species distribution modeling.

We then tested how much niches potentially changed between time periods. To compare the overall difference in niches between each time period, we calculated the multidimensional Euclidean distance between species’ TTR niches for each of the three pairs of time periods. We also compared the change in individual traits by subtracting the values of the earlier niche from the later niche in each pair. We considered all 17 TTR variables selected during data preparation. To test whether levels of niche and trait change were different between time-period pairs, we fitted phylogenetic linear mixed models with time period as the main effect predicting each individual trait difference and Euclidean distance using pglmm as above.

## RESULTS

### Distributions of Veronica through time

The present-day distributions of *Veronica* spanned both temperate and subpolar oceanic climate habitats of New Zealand, with the coldest month averaging above 0°C and with at least four or up to three months, respectively, with mean temperatures above 10°C (Kottek et al., 2006; Rubel & Kottek, 2010). Distributions had shifted marginally since the mid-Holocene and were largely predicted to shrink or retreat southwards by 2070. Across New Zealand, range loss was highest in the most extreme future climate scenario and was higher than range gain, although range retention was generally higher than range loss in the South Island for all scenarios (Fig. S1). The Thornley Transport Resistance model generally fit the observed distributions well for present-day climate. The mean (± standard error) AUC for each species was 0.89 (±0.01). False positives were somewhat inflated with a mean of 0.17 (±0.01) across species but false negatives were much lower at 0.07 (±0.01) (Table S4).

### Range sensitivity to past and future climate change

Ninety-five percent of species were predicted to lose more of their range in the next 50 years than they did since the mid-Holocene, with only twelve species potentially experiencing net range expansion (Fig. 1). The estimated mean percent range loss for species between the mid-Holocene and present climate was 9.0% (±0.6%), which differed from 38.1% (±1.7%) and 54.7% (±2.2%) loss under the moderate and more extreme future climate change scenarios, RCP 4.5 and RCP 8.5, respectively (Fig. 1a, Table S5). Range gain did not differ among scenarios, varying between 10.2% to 18.6%, on average, per species (Table S6), except for seven species that were outliers and gained an estimated mean of 45% to 336% of their present-day range under RCP 8.5 (Fig. 1b). Phylogenetic signal was generally low, suggesting vulnerability was widely dispersed across *Veronica* (Table S7). However, the outliers with the largest increases in range gain all occupied lowland habitats in the North Island and several were closely related (Fig. 1). Range gain was also moderately correlated between past and future scenarios (RCP4.5: r=0.74, RCP8.5: r=0.69), while range loss was weakly correlated (RCP4.5: r=0.45, RCP8.5: r=0.4), suggesting that the underlying factors controlling range gain were more consistent over time (Fig. 2).

**Figure 1.**
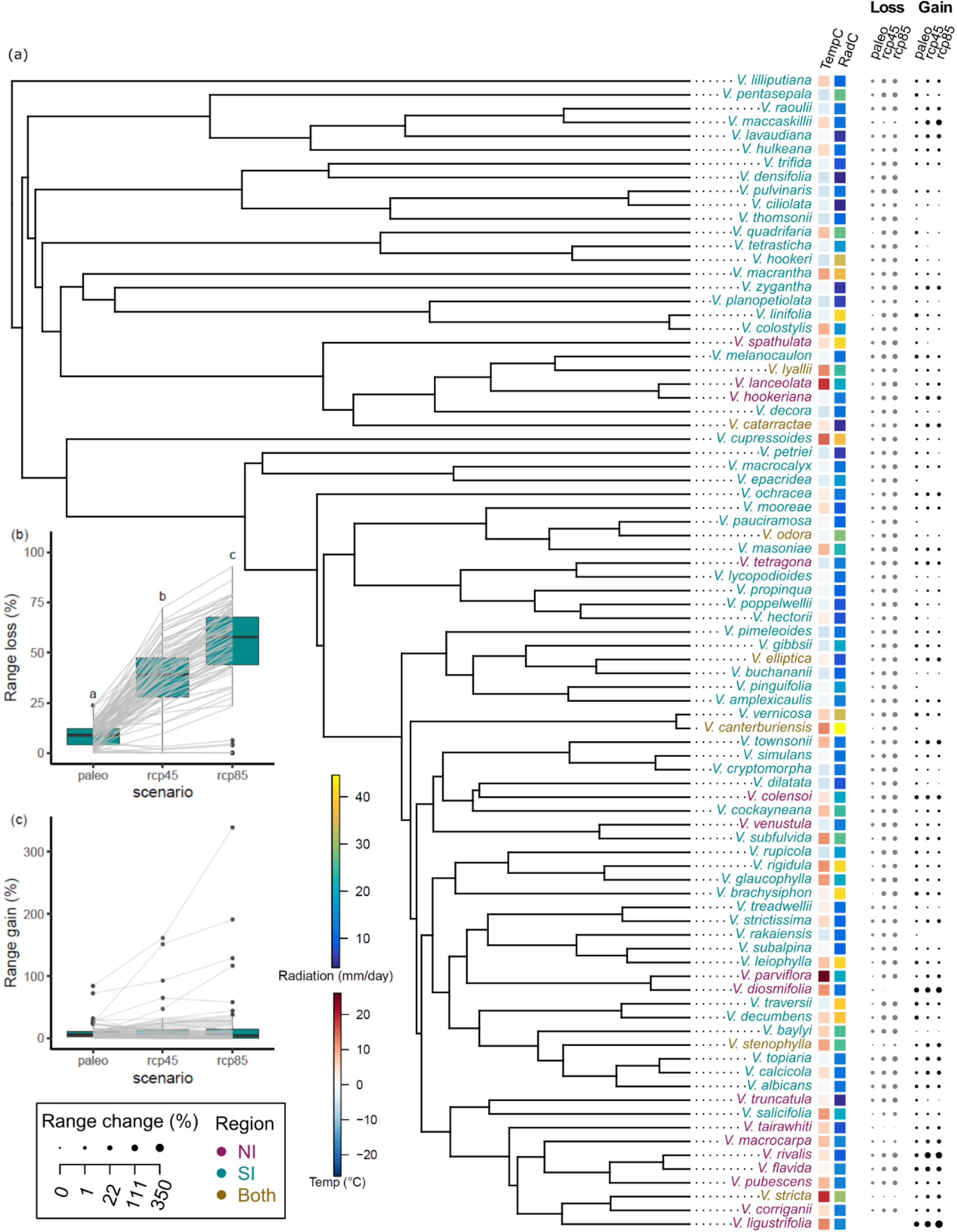
Many losers and a few winners with future climate change in *Veronica*. (**a**) Phylogenetic context for changes in *Veronica*. Circles show size of percent range loss (gray) or gain (black), quarter-root transformed, for each species from mid-Holocene (6000 mya; “paleo”) to current climate (1960-1990) and from current climate to future (2070) scenarios under different atmospheric carbon concentrations (“rcp45” and “rcp85”). Colored squares show species’ saturation thresholds of temperature (TempC) and radiation (RadC) for photosynthesis. For percent of range (**b**) lost and (**c**) gained across time, boxes show median and quartiles; whiskers show minimum and maximum, and black points are outliers that are outside of 1.5-times interquartile range. Gray lines link n=84 species across time periods. Different letters indicate statistically significant differences between scenarios.

**Figure 2.**
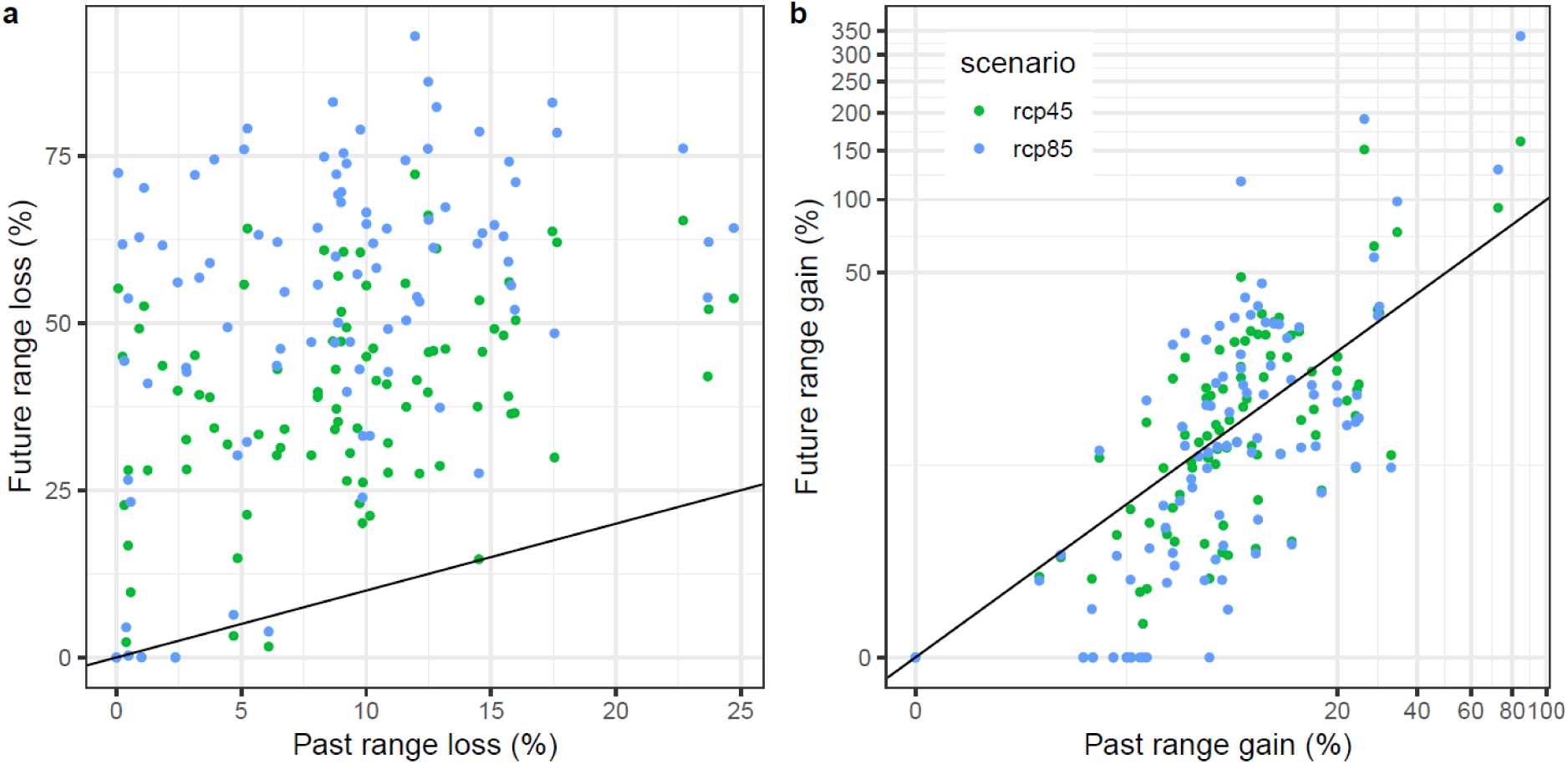
Range gain more strongly correlated between past and future than range loss. Relationships between past range (**a**) loss and (**b**) gain, calculated between mid-Holocene (6000 mya) and current climate (1960-1990), and future loss and gain, calculated between current climate and future (2070) scenarios under different atmospheric carbon concentrations (“rcp45” and “rcp85”). Black lines show 1:1 relationships.

Species’ vulnerability to range loss could be predicted by physiological tolerances and habitat, but only in the future. Over the Holocene period, no traits were associated with increased range loss (Table S5), indicating that the niche tolerances measured here did not strongly influence past range loss. In the future scenarios, however, the most important niche traits for determining range loss were related to solar radiation and temperature associated with photosynthesis (Table S5). Higher radiation thresholds for the saturation of photosynthetic rate were associated with more range loss (Fig. 3a), possibly because species requiring more light would be exposed to hotter, drier conditions under climate change. Over the range of radiation thresholds observed in *Veronica* (1.6 to 20 mm day^-1^ equivalent evapotranspiration), estimated mean range loss increased from 30.1% (95% confidence interval [CI]: 25.5%–34.6%) to 48.3% (39.2%–57.4%) and from 43.2% (38.7%–47.8%) to 68.5% (59.4%–77.6%) under the RCP 4.5 and 8.5 scenarios, respectively, at the mean values of all the other variables. A higher temperature tolerance for photosynthesis had the opposite effect (Fig. 3b). Over the range of temperature saturation thresholds for photosynthesis (-5 to 26°C), estimated range loss decreased from means of 44.2% (38.9%–49.5%) to 13.7% (0%– 28.4%) and 62.7% (57.4 %–68%) to 20.9% (6.3%–35.6%) under the RCP 4.5 and 8.5 scenarios, respectively. Similarly, for the range of upper temperature limits at which photosynthesis declines (2.5 to 40°C), range loss decreased from 42.8% (37.4%–48.1%) to 30.5% (25.9%–35%) and from 57.3% (51.9%–62.6%) to 46.8% (42.3%–51.4%) under the RCP 4.5 and 8.5 scenarios, respectively (Fig. 3c). Finally, mountain species were predicted to lose 1.6- and 1.3-times more range than lowland species under RCP 4.5 and 8.5, respectively (Fig. 4a).

**Figure 3.**
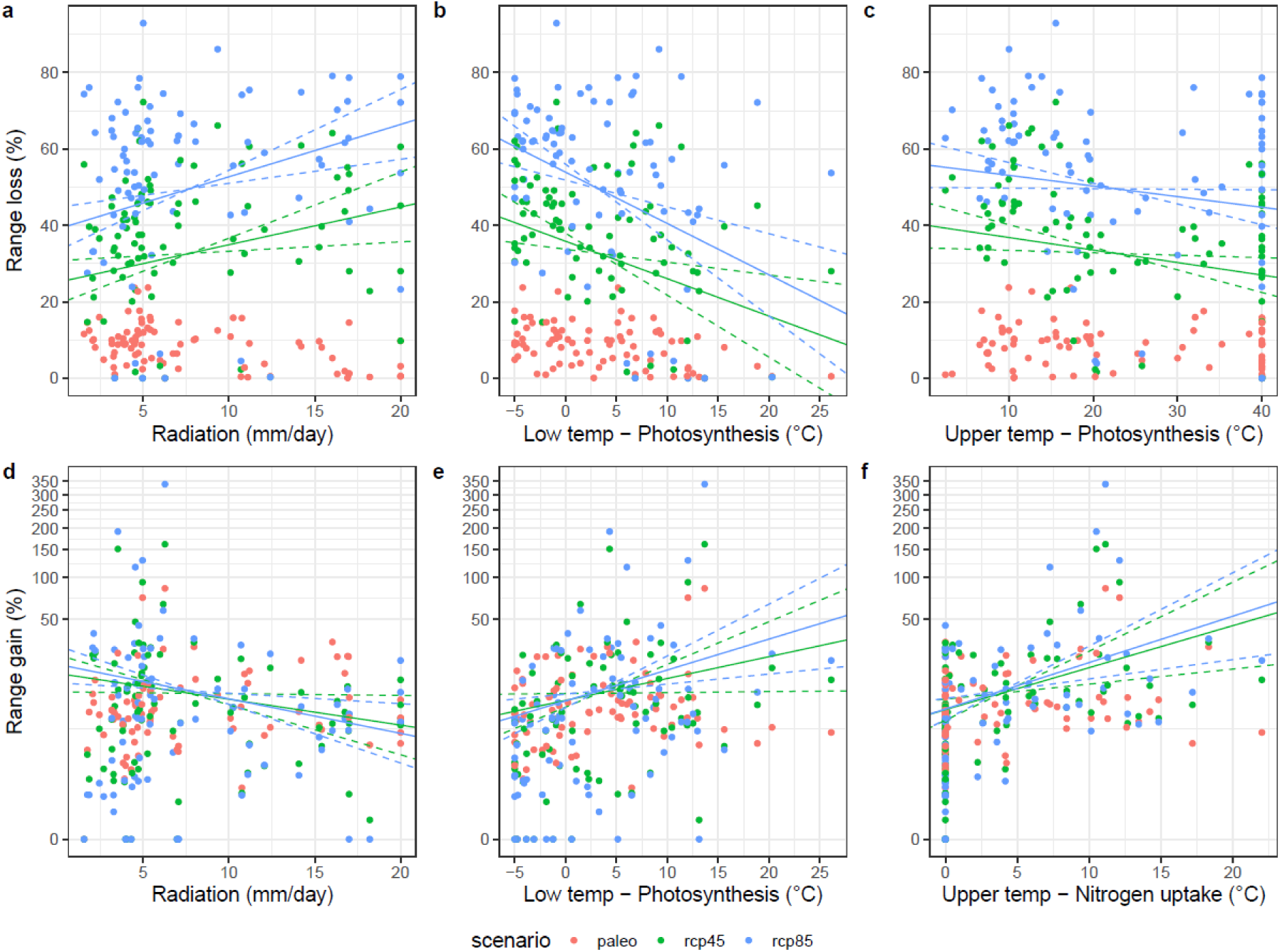
Physiological niche traits predict future range sensitivity to climate change. Relationships between modeled physiological niche traits and (a-c) range loss and (d-f) range gain calculated from mid-Holocene (6000 mya; “paleo”) to current climate (1960-1990) and from current climate to future (2070) scenarios under different atmospheric carbon concentrations (“rcp45” and “rcp85”) for n=84 species. Lines show statistically significant associations predicted by phylogenetic linear mixed models when all other predictors are at their mean values; dashed lines show 95% confidence intervals. Niche traits: (a, d) Radiation threshold at which photosynthesis saturates; (b, e) Lower temperature threshold for saturation of photosynthesis; (c) upper temperature threshold at which photosynthesis declines; (f) saturation threshold of temperature for nitrogen uptake.

**Figure 4.**
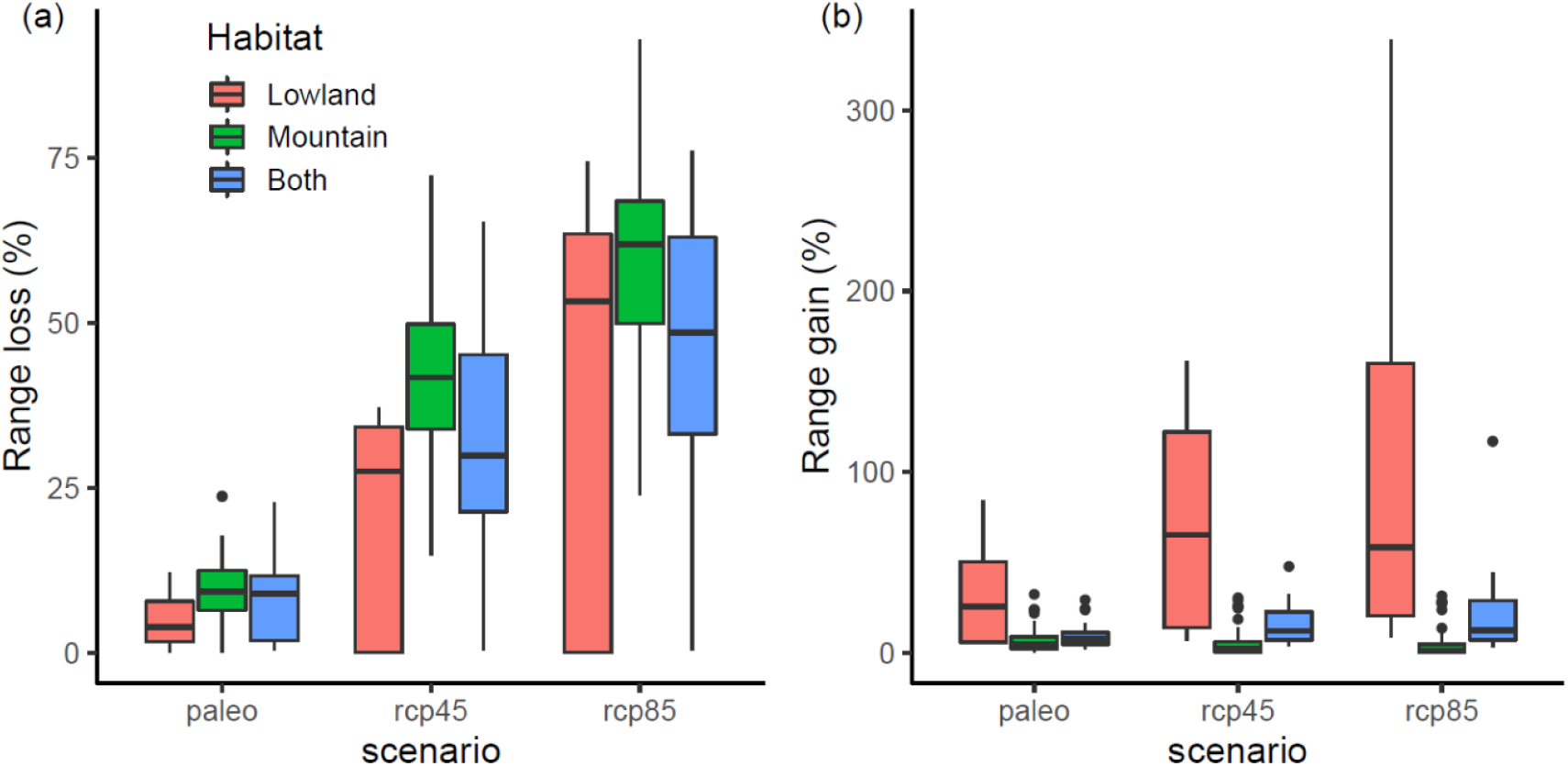
Mountain species more likely to lose range and lowland species more likely to gain range in the future. Predicted percent range (**a**) gain and (**b**) loss from mid-Holocene (6000 mya; “paleo”) to current climate (1960-1990) and from current climate to future (2070) scenarios under different atmospheric carbon concentrations (“rcp45” and “rcp85”). Boxes show median and quartiles; whiskers show minimum and maximum, and black points show outliers >1.5-times the interquartile range. N = 7, 52, and 25 New Zealand *Veronica* species found in lowland, mountain, and both habitats, respectively.

Regardless of time period, lowland species were more likely to gain potential range than mountain species (Fig. 4b). The mean range gain predicted for lowland species was 26.6% (95% CI: 11.4%–53.2%) in the past, which differed from 4.7% (0.9%–15.3%) for mountain species and 6.5% (1.5%–18.7%) for species that occurred in both mountains and lowland (Fig. 2b, Table S6). Lowland range gain was even higher in the future, with similar differences from mountain species. Predicted mean range gain was 30.8% in lowland, 3.1% in mountain specialist, and 6.7% in mountain generalist species under RCP4.5, and 39.3% in lowland, 2.4% in mountain specialist, and 7.0% in mountain generalist species under RCP8.5. Higher shoot nitrogen requirements for photosynthesis also predicted range expansion regardless of time period (Table S6), though the effect was again stronger in the future. Over the range of upper shoot nitrogen concentration thresholds (0.001 to 0.029 kg N per kg tissue), the estimated mean range gain increased from 2.4% (1.1%–4.8%) to 12.6% (6.5%–22.3%) in the past, from 1% (0.3%–2.3%) to 17.7 (9.7%–29.9%) under RCP4.5, and from 0.8% (0.3%–2.3%) to 17.2% (9.4%–29.1%) under RCP 8.5.

Like future range loss, future range gain could also be predicted by physiological tolerances (Table S6). Over the range of radiation thresholds for the saturation of photosynthetic rate, estimated mean range gain decreased from 7.6% (95% CI: 4.8%–11.4%) to 1.4% (0.3%–4.3%) and from 8.2% (5.3%–12.3%) to 0.7% (0.1%–2.7%) under the RCP 4.5 and 8.5 scenarios, respectively (Fig. 3d). By contrast, higher temperature tolerances favored range gain, suggesting sensitivities to radiation may have largely reflected changes in evapotranspiration that are not explicitly modelled in TTR. Over the range of temperature saturation thresholds for photosynthesis, estimated mean range gain increased from 2.4% (1.2%–4.5%) to 18.4% (5%–49.3%) and from 1.6% (0.7%–3.2%) to 27.6% (8.6%–67.8%) under the RCP 4.5 and 8.5 scenarios, respectively (Fig. 3e). Similarly, over the range of temperature saturation thresholds for nitrogen uptake (0 to 22°C), estimated mean range gain increased from 2.6% (1.7%–3.8%) to 32.5% (10.4%–78.8%) and from 2.2% (1.4%–3.3%) to 36.3% (12%–86%) under the RCP 4.5 and 8.5 scenarios, respectively (Fig. 3f). These results suggest that new traits also favor range gain in the future relative to the past, despite similar amounts of gain between periods.

### Niche sensitivity to past and future climate change

To maintain their present-day ranges against future climate change, the temperature tolerances of species will have to change considerably more than they have over the last 6000 years. While there was no predicted change in the temperature tolerances for photosynthesis between the mid-Holocene to the present, the mean upper temperature limits on photosynthesis increased by 3.1 to 3.7 °C (±1.5 °C) between the present day and future scenarios (Table 1). The upper temperature saturation threshold for nitrogen uptake was also predicted to increase by 1.3 to 2.0 °C (±0.6 °C) by 2070 (Table 1). By contrast, species required lower soil moisture content in the present than in the mid-Holocene, with no further change predicted in the future (Table 1). Overall niche distance, measured by the Euclidean distance of 17 modelled physiological traits, did not differ between time periods (Fig. S2), as many of the traits showed a high level of variation among species.

**Table 1.** Different traits influenced range maintenance between the past and future. Mean change (± error) in traits associated with temperature and soil moisture tolerances were compared between mid-Holocene and present climate and between present and future climate scenarios (“rcp45” and “rcp85”). TempC3 is the upper temperature at which photosynthesis begins to decline; TempC4 is the upper temperature limit on photosynthesis (°C); TempN2 is the upper saturation threshold of temperature for nitrogen uptake; SoilmoistN1 and SoilmoistN2 are the lower limit and saturation threshold of percent soil moisture on nitrogen uptake. Colored cells are significantly different from 0 (p<0.05); stars indicate significant difference from the baseline of change between paleo and present (*p<0.05, **p < 0.01).

| | TempC3 ($^{\circ}\text{C}$) | TempC4 ($^{\circ}\text{C}$) | TempN2 ($^{\circ}\text{C}$) | SoilmoistN1 (%) | SoilmoistN2 (%) |
| --- | --- | --- | --- | --- | --- |
| paleo to current | -1.8 ($\pm 1.6$) | -1.1 ($\pm 1.5$) | -0.5 ($\pm 0.6$) | -6.6 ($\pm 3.3$) | -6.6 ($\pm 2.7$) |
| current to rcp45 | 4.0 ($\pm 1.6$)** | 3.1 ($\pm 1.5$)* | 1.3 ($\pm 0.6$)* | 1.9 ($\pm 3.3$)* | 1.3 ($\pm 2.7$)* |
| current to rcp85 | 4.0 ($\pm 1.6$)** | 3.7 ($\pm 1.5$)** | 2.0 ($\pm 0.6$)** | 0.4 ($\pm 3.3$) | 0.3 ($\pm 2.7$)* |

## DISCUSSION

Understanding the historical and physiological context of species’ vulnerabilities to climate change is a crucial step in predicting the impact of climate change on biodiversity (Nadeau et al., 2017). By combining temporally contrasting range and niche projections, we show that range shifts in the next 50 years will need to be more extreme than in the past 6000 years to track climate niches in a large plant radiation. Different physiological niche traits, particularly temperature and radiation tolerance, will be important for predicting range loss and range gain under future compared with past climate change (Antão et al., 2022). In the absence of migration, strong temperature niche shifts will also be required for many species in the future (Jezkova & Wiens, 2016) but not the past. However, some traits remained consistent predictors over time of potential climate winners, notably occupancy of lowland habitat. Our study adds to a growing body of evidence that despite the threats posed by climate change to many species, not all species will experience unmitigated loss, and that it may be possible to predict which species are most at risk based on physiological and geographical traits (Araújo et al., 2011; Dagnino et al., 2020; Thuiller et al., 2005).

Our results demonstrate that the widely acknowledged importance of higher temperature tolerance for promoting future range occupancy (Antão et al., 2022; Feeley et al., 2020; Gottfried et al., 2012; Thuiller et al., 2005) is a more consistent feature of anthropogenic than Holocene climate change. While many plant species did track climate niches to some extent during the Holocene (Corlett & Westcott, 2013; Davis & Shaw, 2001; Liang et al., 2018), indicating that climate tolerances mattered for range occupancy, factors such as precipitation, seasonality, and carbon dioxide concentration may have interacted in different, complex ways with temperature (Davis & Shaw, 2001; Jackson & Overpeck, 2000; Svenning et al., 2015; VanDerWal et al., 2013; Veloz et al., 2012). The unprecedented strength of the future temperature gradient thus likely has an outsized effect on species’ potential ranges relative to the mid-Holocene. At the same time, the negative association between future range size and radiation tolerance shown here suggests a decoupling of light and temperature tolerances. Species that currently favor high light conditions do not necessarily also tolerate higher temperatures, such as the many *Veronica* species that prefer open habitat at high elevations (Bayly & Kellow, 2006), possibly exposing them to future conditions of higher temperature and evapotranspiration than they are adapted to withstand. Thus, both the increased importance of temperature tolerance and no-analog climate combinations will likely be the strongest physiological filters of range occupancy under future climate change (Antão et al., 2022; Gottfried et al., 2012).

Further, when examining potential niche change, we found that species will on average require higher temperature tolerances to occupy the same ranges in the future as in the present day. The average increases in temperature thresholds for photosynthesis and nitrogen uptake were of a similar magnitude to temperature increases projected over a similar period for a range of localities in Jezkova and Wiens (2016), thus roughly tracking climate change. Again, this differed from the Holocene, in which the overall lack of temperature niche change may reflect the slow rates of past niche change modeled by Jezkova and Wiens (2016) over millions of years, as well as the less extreme change in Holocene temperatures. Instead, past niche changes were characterized by an average drop in the lower limit of soil moisture needed for nitrogen uptake, indicating again that axes other than temperature, such as aridity, likely had a stronger relative effect in the past.

Where species track expanding or contracting ranges, our results suggest that communities will be restructured differently in different habitats. Mountain species were more vulnerable to range loss and less likely to gain additional range than lowland species, pointing to loss of community interactions as a dominant future trend in the mountains (Engler et al., 2011; Gottfried et al., 2012). Even with upslope migration, mountain range loss is unsurprising given the limited area of high elevation habitat (Engler et al., 2011; but see Elsen & Tingley, 2015; Liang et al., 2018). However, mountain species’ range contractions could be mitigated to some extent by shifts to microhabitat refugia on mountain slopes not captured by broad-scale climate niche modeling (Graae et al., 2018; Kulonen et al., 2018). In this scenario, with species moving short distances, mountain communities may see less drastic restructuring, at least in the medium term (Hülber et al., 2020). By contrast, lowland range gain tended to occur latitudinally from north to south, requiring species to disperse larger distances to access new habitat. Lowland habitat may thus be more prone to the development of large-scale novel communities and interactions (Antão et al., 2022; Urban et al., 2012). Across habitats, projected range gain was more consistent over time than range loss, suggesting this process may have occurred before. However, the outcome of future range expansion will depend strongly on dispersal capacity, competitive ability, and non-climatic factors in habitat suitability, such as soil type and land use (Bellard et al., 2012; Corlett & Westcott, 2013; Hülber et al., 2020; Spence & Tingley, 2020; Urban et al., 2012).

Our study uses a mechanistic physiological model to overcome some of the limitations of purely correlative SDMs, defining process-based traits that reflect the fundamental niche. This allows us to make niche comparisons less limited by confounding factors controlling species’ realized ranges (Catullo et al., 2015; Higgins et al., 2020). While this approach is useful, complementary approaches are needed to address the full range of factors affecting species’ vulnerability to climate change. As mentioned, species’ dispersal ability and non-climatic tolerances vary widely and will determine access to climatically suitable range. In addition, adaptive potential is shaped by past variability in climate in ways that snapshots of past and current climate alone cannot capture, calling for studies including longer term climate trends (Nadeau et al., 2017). Finally, we examined potential range shifts and niche shifts separately, but in reality, both will likely occur together (Davis & Shaw, 2001; Lustenhouwer & Parker, 2022). A moderate level of niche evolution or plasticity could reduce the distance needed to migrate to track climate, increase dispersal capacity (Nadeau & Urban, 2019), or mitigate mismatches of environmental axes to open up unforeseen range availability (Lustenhouwer et al., 2018). A key role of future studies will be to develop methods that can combine predictions of potential range and niche change, for example by using models that take into account both dispersal (e.g. Engler et al., 2009) and the possibility of niche adaptation (Catullo et al., 2015; Hua et al., 2021) in both past and future. Continuing to develop the ability of niche models to inform conservation planning for vulnerable species is vital in the fight against biodiversity loss with climate change.

