## Supplementary tables and figures for "Winners and losers under past and future climate change"

**Table S1 | Selected TTR niche traits with illustrations of their physiological responses along environmental axes.** Response functions for each variable are either trapezoidal or stepwise. Trapezoidal variables have four parameters delineating at what values the process: (i) begins to increase from zero, (ii) saturates, that is, further increases in the environmental parameter do not increase the physiological response (labelled 1 on the y-axis), (iii) is once again reduced, and (iv) can no longer occur. Two-step functions have (i) a lower limit at which the process begins to increase from zero and (ii) a value above which the process is saturated. Grey dots designate selected parameter from the trapezoid or two-step function. *denotes traits included in phylogenetic linear mixed models (Tables S5, S6)

| Parameter | Trait | Function |
|---|---|---|
| TempC1 | Lower temperature limit for carbon uptake (photosynthesis) | 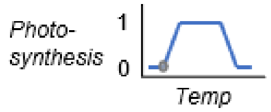  |
| TempC2* | Lower temperature saturation threshold at which carbon uptake no longer increases with higher temps | 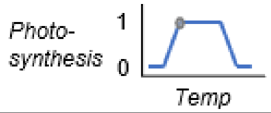  |
| TempC3* | Upper temperature limit at which carbon uptake begins to decrease | 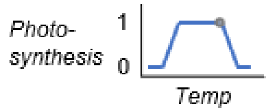  |
| TempC4 | Upper temperature limit for carbon uptake | 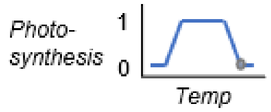  |
| RadC2* | Solar radiation saturation threshold at which carbon uptake no longer increases with higher radiation | 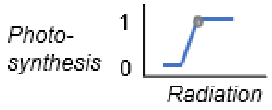  |
| SoilmoistC1* | Lower limit of soil moisture for carbon uptake | 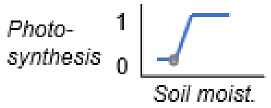  |
| NshootC2* | Shoot nitrogen saturation threshold for carbon uptake | 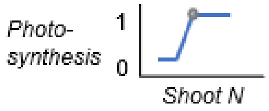  |
| TempN2* | Temperature saturation threshold for nitrogen uptake | 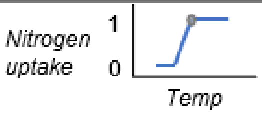  |
| SoilmoistN1* | Lower limit of soil moisture for nitrogen uptake | 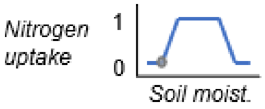  |
| SoilmoistN2 | Soil moisture saturation threshold for nitrogen uptake | 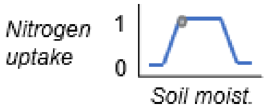  |
| TempResp2* | Temperature saturation threshold for respiration | 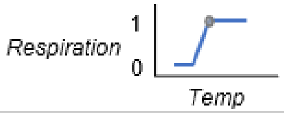  |

**Table S2 | *Veronica* species included in the analysis and details of TTR performance.** Habitat is mountain (M), lowland (L), or both (ML). TP=true positives of TTR fit on testing data; TN=true negatives; FP=false positives; FN=false negatives; AUC=area under the receiver-operator curve; AUC.sd=standard deviation of AUC.

| Species | Habitat | Total occ | TP | TN | FP | FN | AUC | AUC.sd |
|---|---|---|---|---|---|---|---|---|
| **V. albicans** | ML | 140 | 0.94 | 0.88 | 0.12 | 0.06 | 0.95 | 0.03 |
| **V. amplexicaulis** | M | 37 | 0.9 | 0.78 | 0.22 | 0.1 | 0.85 | 0.09 |
| **V. baylyi** | M | 29 | 0.88 | 0.86 | 0.14 | 0.12 | 0.87 | 0.09 |
| **V. brachysiphon** | M | 104 | 0.88 | 0.84 | 0.16 | 0.12 | 0.89 | 0.04 |
| **V. buchananii** | M | 111 | 1 | 0.78 | 0.22 | 0 | 0.92 | 0.03 |
| **V. calcicola** | M | 27 | 1 | 0.75 | 0.25 | 0 | 0.79 | 0.12 |
| **V. canterburiensis** | M | 181 | 1 | 0.87 | 0.13 | 0 | 0.98 | 0.02 |
| **V. catarractae** | ML | 80 | 0.94 | 0.89 | 0.11 | 0.06 | 0.93 | 0.05 |
| **V. ciliolata** | M | 74 | 1 | 0.95 | 0.05 | 0 | 0.96 | 0.03 |
| **V. cockayneana** | M | 74 | 0.95 | 0.9 | 0.1 | 0.05 | 0.93 | 0.05 |
| **V. colensoi** | ML | 57 | 1 | 0.77 | 0.23 | 0 | 0.99 | 0.01 |
| **V. colostylis** | M | 61 | 0.81 | 0.78 | 0.22 | 0.19 | 0.91 | 0.06 |
| **V. corriganii** | ML | 77 | 1 | 0.65 | 0.35 | 0 | 0.92 | 0.04 |
| **V. cryptomorpha** | M | 36 | 0.78 | 0.8 | 0.2 | 0.22 | 0.96 | 0.04 |
| **V. cupressoides** | M | 51 | 0.85 | 0.58 | 0.42 | 0.15 | 0.84 | 0.08 |
| **V. decora** | M | 190 | 0.91 | 0.64 | 0.36 | 0.09 | 0.78 | 0.05 |
| **V. decumbens** | M | 80 | 0.95 | 0.89 | 0.11 | 0.05 | 0.9 | 0.05 |
| **V. densifolia** | M | 82 | 0.95 | 0.87 | 0.13 | 0.05 | 0.89 | 0.05 |
| **V. dilatata** | M | 26 | 0.86 | 1 | 0 | 0.14 | 0.93 | 0.07 |
| **V. diosmifolia** | L | 74 | 0.94 | 0.88 | 0.12 | 0.06 | 0.93 | 0.05 |
| **V. elliptica** | L | 68 | 0.92 | 0.41 | 0.59 | 0.08 | 0.56 | 0.11 |
| **V. epacridea** | M | 169 | 0.93 | 0.9 | 0.1 | 0.07 | 0.88 | 0.04 |
| **V. flavida** | L | 28 | 1 | 1 | 0 | 0 | 1 | 0 |
| **V. gibbsii** | M | 20 | 1 | 0.8 | 0.2 | 0 | 0.88 | 0.12 |
| **V. glaucophylla** | M | 52 | 0.69 | 0.8 | 0.2 | 0.31 | 0.67 | 0.11 |
| **V. hectorii** | M | 300 | 0.93 | 0.77 | 0.23 | 0.07 | 0.79 | 0.04 |

| | | | | | | | | |
|---|---|---|---|---|---|---|---|---|
| **V. hookeri** | M | 173 | 0.91 | 0.86 | 0.14 | 0.09 | 0.93 | 0.03 |
| **V. hookeriana** | M | 117 | 0.87 | 0.89 | 0.11 | 0.13 | 0.85 | 0.05 |
| **V. hulkeana** | ML | 64 | 0.88 | 0.88 | 0.12 | 0.12 | 0.97 | 0.02 |
| **V. lanceolata** | ML | 419 | 0.98 | 0.82 | 0.18 | 0.02 | 0.89 | 0.03 |
| **V. lavaudiana** | ML | 35 | 0.88 | 0.8 | 0.2 | 0.12 | 0.78 | 0.1 |
| **V. leiophylla** | ML | 164 | 0.85 | 0.74 | 0.26 | 0.15 | 0.85 | 0.04 |
| **V. ligustrifolia** | L | 89 | 1 | 0.96 | 0.04 | 0 | 0.99 | 0.01 |
| **V. lilliputiana** | ML | 59 | 0.73 | 0.93 | 0.07 | 0.27 | 0.75 | 0.09 |
| **V. linifolia** | M | 104 | 0.96 | 0.85 | 0.15 | 0.04 | 0.92 | 0.04 |
| **V. lyallii** | ML | 494 | 0.84 | 0.66 | 0.34 | 0.16 | 0.85 | 0.02 |
| **V. lycopodioides** | M | 178 | 0.98 | 0.73 | 0.27 | 0.02 | 0.86 | 0.04 |
| **V. maccaskillii** | ML | 27 | 1 | 0.67 | 0.33 | 0 | 0.76 | 0.13 |
| **V. macrantha** | M | 154 | 0.84 | 0.9 | 0.1 | 0.16 | 0.93 | 0.03 |
| **V. macrocalyx** | M | 117 | 0.97 | 0.83 | 0.17 | 0.03 | 0.89 | 0.04 |
| **V. macrocarpa** | ML | 160 | 0.94 | 0.91 | 0.09 | 0.06 | 0.95 | 0.03 |
| **V. masoniae** | M | 60 | 0.87 | 0.93 | 0.07 | 0.13 | 0.89 | 0.06 |
| **V. melanocaulon** | ML | 53 | 1 | 0.93 | 0.07 | 0 | 0.92 | 0.06 |
| **V. mooreae** | M | 128 | 0.97 | 0.81 | 0.19 | 0.03 | 0.92 | 0.03 |
| **V. ochracea** | M | 37 | 0.9 | 0.9 | 0.1 | 0.1 | 0.95 | 0.05 |
| **V. odora** | M | 429 | 0.8 | 0.84 | 0.16 | 0.2 | 0.83 | 0.03 |
| **V. parviflora** | ML | 147 | 1 | 0.54 | 0.46 | 0 | 0.74 | 0.06 |
| **V. pauciramosa** | M | 140 | 0.91 | 0.88 | 0.12 | 0.09 | 0.89 | 0.04 |
| **V. pentasepala** | M | 32 | 1 | 1 | 0 | 0 | 1 | 0 |
| **V. petriei** | M | 30 | 0.75 | 1 | 0 | 0.25 | 0.94 | 0.06 |
| **V. pimeleoides** | M | 111 | 0.93 | 0.57 | 0.43 | 0.07 | 0.76 | 0.06 |
| **V. pinguifolia** | M | 183 | 0.91 | 0.71 | 0.29 | 0.09 | 0.82 | 0.05 |
| **V. planopetiolata** | M | 27 | 1 | 0.88 | 0.12 | 0 | 1 | 0 |
| **V. poppelwellii** | M | 31 | 1 | 1 | 0 | 0 | 0.69 | 0.09 |
| **V. propinqua** | M | 64 | 1 | 0.44 | 0.56 | 0 | 0.85 | 0.07 |
| **V. pubescens** | L | 54 | 0.91 | 0.86 | 0.14 | 0.09 | 0.87 | 0.08 |
| **V. pulvinaris** | M | 146 | 0.89 | 0.91 | 0.09 | 0.11 | 0.84 | 0.05 |

| | | | | | | | | |
|---|---|---|---|---|---|---|---|---|
| **V. quadrifaria** | M | 70 | 0.94 | 0.88 | 0.12 | 0.06 | 0.89 | 0.05 |
| **V. rakaiensis** | M | 99 | 0.96 | 0.64 | 0.36 | 0.04 | 0.87 | 0.05 |
| **V. raoulii** | ML | 75 | 0.74 | 0.78 | 0.22 | 0.26 | 0.82 | 0.07 |
| **V. rigidula** | ML | 45 | 1 | 0.77 | 0.23 | 0 | 0.9 | 0.06 |
| **V. rivalis** | L | 36 | 1 | 0.89 | 0.11 | 0 | 1 | 0 |
| **V. rupicola** | ML | 39 | 1 | 0.9 | 0.1 | 0 | 0.8 | 0.1 |
| **V. salicifolia** | ML | 243 | 0.96 | 0.63 | 0.37 | 0.04 | 0.93 | 0.02 |
| **V. simulans** | M | 60 | 1 | 0.86 | 0.14 | 0 | 0.98 | 0.02 |
| **V. spathulata** | M | 62 | 1 | 0.94 | 0.06 | 0 | 0.93 | 0.05 |
| **V. stenophylla** | ML | 190 | 0.89 | 0.66 | 0.34 | 0.11 | 0.85 | 0.04 |
| **V. stricta** | ML | 643 | 0.98 | 0.45 | 0.55 | 0.02 | 0.84 | 0.02 |
| **V. strictissima** | ML | 51 | 1 | 0.8 | 0.2 | 0 | 0.92 | 0.06 |
| **V. subalpina** | M | 241 | 0.98 | 0.82 | 0.18 | 0.02 | 0.95 | 0.02 |
| **V. subfulvida** | ML | 82 | 1 | 0.59 | 0.41 | 0 | 0.8 | 0.07 |
| **V. tairawhiti** | L | 22 | 1 | 1 | 0 | 0 | 1 | 0 |
| **V. tetragona** | M | 141 | 0.8 | 0.94 | 0.06 | 0.2 | 0.85 | 0.04 |
| **V. tetrasticha** | M | 50 | 0.85 | 1 | 0 | 0.15 | 1 | 0 |
| **V. thomsonii** | M | 79 | 0.74 | 1 | 0 | 0.26 | 0.88 | 0.05 |
| **V. topiaria** | M | 99 | 0.92 | 0.81 | 0.19 | 0.08 | 0.9 | 0.05 |
| **V. townsonii** | ML | 20 | 1 | 1 | 0 | 0 | 1 | 0 |
| **V. traversii** | ML | 161 | 0.93 | 0.78 | 0.22 | 0.07 | 0.95 | 0.03 |
| **V. treadwellii** | M | 55 | 1 | 0.73 | 0.27 | 0 | 0.9 | 0.06 |
| **V. trifida** | M | 40 | 1 | 0.9 | 0.1 | 0 | 0.99 | 0.01 |
| **V. truncatula** | M | 22 | 1 | 0.86 | 0.14 | 0 | 0.98 | 0.03 |
| **V. venustula** | ML | 97 | 0.72 | 0.95 | 0.05 | 0.28 | 0.87 | 0.05 |
| **V. vernicosa** | M | 118 | 0.93 | 0.74 | 0.26 | 0.07 | 0.93 | 0.03 |
| **V. zygantha** | M | 22 | 1 | 1 | 0 | 0 | 0.91 | 0.09 |

**Table S3 | Phylogenetic principal components analysis loadings** of TTR traits used to select trait subset for phylogenetic linear mixed modeling. Loadings were examined in the first 7 principal component axes (PCs), which cumulatively explained 95% of the variance (individual PC weights are listed in the final row of the table). Trait loadings were excluded (set to 0) from a given PC if they were below the expected value for single variable's proportion of the sum of squares, 1/sqrt(n) where n=17, the total number of traits in the PCA. Loadings were then weighted by the variance explained by the PC and summed. Where traits were correlated in biplots, the trait with the higher loading score was included in the model.

| Trait | PC1 | PC2 | PC3 | PC4 | PC5 | PC6 | PC7 | Variable weight | Correlate exclude | Include |
|---|---|---|---|---|---|---|---|---|---|---|
| **NshootC1** | 0 | 0 | 0 | 0 | 0 | 0.32 | 0.33 | 0.02 | TRUE | FALSE |
| **NshootC2** | 0 | 0 | 0 | 0 | 0 | 0.47 | 0.71 | 0.03 | FALSE | TRUE |
| **NsoilN1** | 0 | 0 | 0 | 0 | 0 | 0 | 0 | 0 | FALSE | FALSE |
| **NsoilN2** | 0 | 0 | 0 | 0 | 0 | 0 | 0 | 0 | FALSE | FALSE |
| **RadC1** | 0.35 | 0 | 0.35 | 0.36 | 0 | 0 | 0 | 0.21 | TRUE | FALSE |
| **RadC2** | 0.35 | 0 | 0.39 | 0.39 | 0 | 0 | 0 | 0.22 | FALSE | TRUE |
| **TempC1** | 0 | 0 | 0 | 0 | 0 | 0.38 | 0.44 | 0.02 | TRUE | FALSE |
| **TempC2** | 0 | 0 | 0.25 | 0 | 0 | 0.42 | 0.38 | 0.06 | FALSE | TRUE |
| **TempC3** | 0.30 | 0 | 0.50 | 0.36 | 0 | 0 | 0 | 0.22 | FALSE | TRUE |
| **TempC4** | 0.25 | 0 | 0.51 | 0.36 | 0 | 0 | 0 | 0.21 | TRUE | FALSE |
| **TempN2** | 0 | 0 | 0 | 0 | 0 | 0.46 | 0 | 0.01 | FALSE | TRUE |
| **TempR1** | 0 | 0 | 0 | 0 | 0.63 | 0 | 0 | 0.06 | TRUE | FALSE |
| **TempR2** | 0 | 0.24 | 0 | 0 | 0.66 | 0 | 0 | 0.11 | FALSE | TRUE |
| **SoilMoistC1** | 0.36 | 0.39 | 0 | 0.45 | 0 | 0 | 0 | 0.25 | FALSE | TRUE |
| **SoilMoistC2** | 0.32 | 0.38 | 0 | 0.39 | 0 | 0 | 0 | 0.23 | TRUE | FALSE |
| **SoilMoistN1** | 0.44 | 0.50 | 0 | 0 | 0 | 0 | 0 | 0.24 | FALSE | TRUE |
| **SoilMoistN2** | 0.38 | 0.47 | 0 | 0 | 0 | 0 | 0 | 0.21 | TRUE | FALSE |
| *PC weight* | 0.31 | 0.21 | 0.16 | 0.13 | 0.10 | 0.03 | 0.02 | | | |

**Table S4 | Summary of model assessment metrics for TTR models** fit to present day distribution under different climate scenarios, including Area Under the Curve (AUC), false negative rate (FN), and false positive rate (FP).

| Scenario | Metric | Mean | SE |
|---|---|---|---|
| current | AUC | 0.89 | 0.009 |
| | FN | 0.07 | 0.009 |
| | FP | 0.17 | 0.014 |
| paleo | AUC | 0.88 | 0.009 |
| | FN | 0.07 | 0.008 |
| | FP | 0.17 | 0.014 |
| rcp45 | AUC | 0.88 | 0.010 |
| | FN | 0.06 | 0.008 |
| | FP | 0.18 | 0.015 |
| rcp85 | AUC | 0.88 | 0.010 |
| | FN | 0.07 | 0.011 |
| | FP | 0.20 | 0.016 |

**Table S5 | Phylogenetic linear mixed models of selected niche traits and proportion range loss between periods**: paleo (6000 mya) to present (1960-1990 mean) climate, and present to future (2070) scenarios of carbon concentration pathways (RCP4.5 and RCP8.5), shown with each scenario as the baseline as labelled from left to right. For effect means, purple cells show a negative relationship; yellow cells show positive relationship; gray cells are not statistically significant. Bold text marks significant effects and confidence intervals.

| Percent loss ~ traits | | | | | | | | | | | |
|---|---|---|---|---|---|---|---|---|---|---|---|
| Paleo | | | | RCP45 | | | | RCP85 | | | |
| Variable | Effect | Lower.CI | Upper.CI | Variable | Effect | Lower.CI | Upper.CI | Variable | Effect | Lower.CI | Upper.CI |
| (Intercept) | 0.042 | -0.069 | 0.153 | (Intercept) | **0.256** | **0.146** | **0.367** | (Intercept) | **0.453** | **0.343** | **0.564** |
| Partial effects | | | | | | | | | | | |
| RCP45 | **0.215** | **0.109** | **0.320** | Paleo | **-0.215** | **-0.320** | **-0.109** | Paleo | **-0.411** | **-0.516** | **-0.306** |
| RCP85 | **0.411** | **0.306** | **0.516** | RCP85 | **0.197** | **0.092** | **0.302** | RCP45 | **-0.197** | **-0.302** | **-0.092** |
| NshootC2 | 0.007 | -0.025 | 0.038 | NshootC2 | 0.026 | -0.006 | 0.057 | NshootC2 | 0.020 | -0.011 | 0.051 |
| RadC2 | -0.022 | -0.061 | 0.016 | RadC2 | **0.052** | **0.013** | **0.091** | RadC2 | **0.072** | **0.033** | **0.111** |
| TmaxC2 | -0.006 | -0.050 | 0.037 | TmaxC2 | **-0.066** | **-0.110** | **-0.023** | TmaxC2 | **-0.091** | **-0.134** | **-0.047** |
| TmaxC3 | -0.004 | -0.038 | 0.030 | TmaxC3 | **-0.042** | **-0.076** | **-0.008** | TmaxC3 | **-0.036** | **-0.070** | **-0.002** |
| TmeanN2 | 0.008 | -0.036 | 0.052 | TmeanN2 | 0.037 | -0.008 | 0.081 | TmeanN2 | 0.043 | -0.001 | 0.088 |
| TmeanR2 | 0.003 | -0.028 | 0.033 | TmeanR2 | 0.012 | -0.018 | 0.043 | TmeanR2 | 0.010 | -0.020 | 0.041 |
| SoilmoistC1 | 0.004 | -0.027 | 0.035 | SoilmoistC1 | -0.010 | -0.041 | 0.021 | SoilmoistC1 | -0.003 | -0.033 | 0.028 |
| SoilmoistN1 | 0.006 | -0.026 | 0.039 | SoilmoistN1 | -0.004 | -0.036 | 0.028 | SoilmoistN1 | 0.008 | -0.025 | 0.040 |
| HabitatM | 0.053 | -0.075 | 0.182 | HabitatM | **0.166** | **0.038** | **0.295** | HabitatM | **0.140** | **0.011** | **0.268** |
| HabitatML | 0.040 | -0.083 | 0.163 | HabitatML | 0.044 | -0.079 | 0.166 | HabitatML | -0.013 | -0.136 | 0.110 |
| Interactions | | | | | | | | | | | |
| RCP45:NshootC2 | 0.019 | -0.010 | 0.048 | Paleo:NshootC2 | -0.019 | -0.048 | 0.010 | Paleo:NshootC2 | -0.014 | -0.043 | 0.016 |
| RCP85:NshootC2 | 0.014 | -0.016 | 0.043 | RCP85:NshootC2 | -0.006 | -0.035 | 0.024 | RCP45:NshootC2 | 0.006 | -0.024 | 0.035 |
| RCP45:RadC2 | **0.074** | **0.038** | **0.111** | Paleo:RadC2 | **-0.074** | **-0.111** | **-0.038** | Paleo:RadC2 | **-0.094** | **-0.131** | **-0.058** |

| Percent loss ~ traits | | | | | | | | | | | |
|---|---|---|---|---|---|---|---|---|---|---|---|
| Paleo | | | | RCP45 | | | | RCP85 | | | |
| Variable | Effect | Lower.CI | Upper.CI | Variable | Effect | Lower.CI | Upper.CI | Variable | Effect | Lower.CI | Upper.CI |
| RCP85:RadC2 | **0.094** | **0.058** | **0.131** | RCP85:RadC2 | 0.020 | -0.016 | 0.056 | RCP45:RadC2 | -0.020 | -0.056 | 0.016 |
| RCP45:TmaxC2 | **-0.060** | **-0.101** | **-0.019** | Paleo:TmaxC2 | **0.060** | **0.019** | **0.101** | Paleo:TmaxC2 | **0.085** | **0.044** | **0.125** |
| RCP85:TmaxC2 | **-0.085** | **-0.125** | **-0.044** | RCP85:TmaxC2 | -0.025 | -0.065 | 0.016 | RCP45:TmaxC2 | 0.025 | -0.016 | 0.065 |
| RCP45:TmaxC3 | **-0.039** | **-0.070** | **-0.007** | Paleo:TmaxC3 | **0.039** | **0.007** | **0.070** | Paleo:TmaxC3 | **0.032** | **0.001** | **0.064** |
| RCP85:TmaxC3 | **-0.032** | **-0.064** | **-0.001** | RCP85:TmaxC3 | 0.006 | -0.025 | 0.038 | RCP45:TmaxC3 | -0.006 | -0.038 | 0.025 |
| RCP45:TmeanN2 | 0.028 | -0.013 | 0.070 | Paleo:TmeanN2 | -0.028 | -0.070 | 0.013 | Paleo:TmeanN2 | -0.035 | -0.077 | 0.006 |
| RCP85:TmeanN2 | 0.035 | -0.006 | 0.077 | RCP85:TmeanN2 | 0.007 | -0.035 | 0.048 | RCP45:TmeanN2 | -0.007 | -0.048 | 0.035 |
| RCP45:TmeanR2 | 0.010 | -0.019 | 0.038 | Paleo:TmeanR2 | -0.010 | -0.038 | 0.019 | Paleo:TmeanR2 | -0.008 | -0.036 | 0.021 |
| RCP85:TmeanR2 | 0.008 | -0.021 | 0.036 | RCP85:TmeanR2 | -0.002 | -0.030 | 0.027 | RCP45:TmeanR2 | 0.002 | -0.027 | 0.030 |
| RCP45:SoilmoistC1 | -0.014 | -0.043 | 0.015 | Paleo:SoilmoistC1 | 0.014 | -0.015 | 0.043 | Paleo:SoilmoistC1 | 0.006 | -0.023 | 0.035 |
| RCP85:SoilmoistC1 | -0.006 | -0.035 | 0.023 | RCP85:SoilmoistC1 | 0.008 | -0.021 | 0.036 | RCP45:SoilmoistC1 | -0.008 | -0.036 | 0.021 |
| RCP45:SoilmoistN1 | -0.010 | -0.041 | 0.020 | Paleo:SoilmoistN1 | 0.010 | -0.020 | 0.041 | Paleo:SoilmoistN1 | -0.001 | -0.031 | 0.029 |
| RCP85:SoilmoistN1 | 0.001 | -0.029 | 0.031 | RCP85:SoilmoistN1 | 0.012 | -0.018 | 0.042 | RCP45:SoilmoistN1 | -0.012 | -0.042 | 0.018 |
| RCP45:HabitatM | 0.113 | -0.007 | 0.233 | Paleo:HabitatM | -0.113 | -0.233 | 0.007 | Paleo:HabitatM | -0.086 | -0.206 | 0.033 |
| RCP85:HabitatM | 0.086 | -0.033 | 0.206 | RCP85:HabitatM | -0.027 | -0.146 | 0.093 | RCP45:HabitatM | 0.027 | -0.093 | 0.146 |
| RCP45:HabitatML | 0.003 | -0.111 | 0.118 | Paleo:HabitatML | -0.003 | -0.118 | 0.111 | Paleo:HabitatML | 0.053 | -0.061 | 0.168 |
| RCP85:HabitatML | -0.053 | -0.168 | 0.061 | RCP85:HabitatML | -0.057 | -0.171 | 0.058 | RCP45:HabitatML | 0.057 | -0.058 | 0.171 |

**Table S6 | Phylogenetic linear mixed models of selected niche traits and percent range gain (quarter-root transformed) between periods**: paleo (6000 mya) to present (1960-1990 mean) climate, and present to future (2070) scenarios of carbon concentration pathways (RCP4.5 and RCP8.5), shown with each scenario as the baseline as labelled from left to right. For effect means, purple cells show a negative relationship; yellow cells show positive relationship; gray cells are not statistically significant. Bold text marks significant effects and confidence intervals.

| Percent gain ~ traits | | | | | | | | | | | |
|---|---|---|---|---|---|---|---|---|---|---|---|
| Paleo | | | | RCP45 | | | | RCP85 | | | |
| Variable | Effect | Lower. CI | Upper. CI | Variable | Effect | Lower. CI | Upper. CI | Variable | Effect | Lower. CI | Upper. CI |
| (Intercept) | **0.718** | **0.581** | **0.854** | (Intercept) | **0.745** | **0.608** | **0.881** | (Intercept) | **0.792** | **0.655** | **0.928** |
| Partial effects | | | | | | | | | | | |
| RCP45 | 0.027 | -0.073 | 0.128 | Paleo | -0.027 | -0.128 | 0.073 | Paleo | -0.074 | -0.174 | 0.027 |
| RCP85 | 0.074 | -0.027 | 0.174 | RCP85 | 0.047 | -0.054 | 0.147 | RCP45 | -0.047 | -0.147 | 0.054 |
| NshootC2 | **0.047** | **0.009** | **0.086** | NshootC2 | **0.079** | **0.040** | **0.118** | NshootC2 | **0.081** | **0.042** | **0.120** |
| RadC2 | 0.027 | -0.021 | 0.076 | RadC2 | **-0.052** | **-0.100** | **-0.004** | RadC2 | **-0.069** | **-0.118** | **-0.021** |
| TmaxC2 | -0.005 | -0.059 | 0.049 | TmaxC2 | **0.057** | **0.002** | **0.111** | TmaxC2 | **0.080** | **0.026** | **0.134** |
| TmaxC3 | -0.030 | -0.072 | 0.012 | TmaxC3 | -0.026 | -0.069 | 0.016 | TmaxC3 | -0.030 | -0.072 | 0.012 |
| TmeanN2 | 0.028 | -0.027 | 0.083 | TmeanN2 | **0.086** | **0.031** | **0.141** | TmeanN2 | **0.095** | **0.040** | **0.150** |
| TmeanR2 | -0.001 | -0.039 | 0.036 | TmeanR2 | 0.001 | -0.037 | 0.039 | TmeanR2 | 0.008 | -0.030 | 0.046 |
| SoilmoistC1 | 0.007 | -0.032 | 0.045 | SoilmoistC1 | 0.008 | -0.031 | 0.046 | SoilmoistC1 | 0.014 | -0.024 | 0.053 |
| SoilmoistN1 | 0.005 | -0.035 | 0.045 | SoilmoistN1 | 0.021 | -0.019 | 0.061 | SoilmoistN1 | 0.007 | -0.034 | 0.047 |
| HabitatM | **-0.252** | **-0.411** | **-0.092** | HabitatM | **-0.325** | **-0.484** | **-0.165** | HabitatM | **-0.396** | **-0.556** | **-0.237** |
| HabitatML | **-0.213** | **-0.365** | **-0.060** | HabitatML | **-0.237** | **-0.389** | **-0.084** | HabitatML | **-0.277** | **-0.430** | **-0.125** |
| Interactions | | | | | | | | | | | |
| RCP45:NshootC2 | **0.032** | **0.004** | **0.060** | Paleo:NshootC2 | **-0.032** | **-0.060** | **-0.004** | Paleo:NshootC2 | **-0.034** | **-0.062** | **-0.006** |
| RCP85:NshootC2 | **0.034** | **0.006** | **0.062** | RCP85:NshootC2 | 0.002 | -0.026 | 0.030 | RCP45:NshootC2 | -0.002 | -0.030 | 0.026 |
| RCP45:RadC2 | **-0.079** | **-0.114** | **-0.045** | Paleo:RadC2 | **0.079** | **0.045** | **0.114** | Paleo:RadC2 | **0.097** | **0.062** | **0.132** |
| RCP85:RadC2 | **-0.097** | **-0.132** | **-0.062** | RCP85:RadC2 | -0.017 | -0.052 | 0.017 | RCP45:RadC2 | 0.017 | -0.017 | 0.052 |
| RCP45:TmaxC2 | **0.061** | **0.022** | **0.100** | Paleo:TmaxC2 | **-0.061** | **-0.100** | **-0.022** | Paleo:TmaxC2 | **-0.085** | **-0.123** | **-0.046** |

### Percent gain ~ traits

| Paleo | | | | RCP45 | | | | RCP85 | | | |
|---|---|---|---|---|---|---|---|---|---|---|---|
| Variable | Effect | Lower. CI | Upper. CI | Variable | Effect | Lower. CI | Upper. CI | Variable | Effect | Lower. CI | Upper. CI |
| RCP85:TmaxC2 | **0.085** | **0.046** | **0.123** | RCP85:TmaxC2 | 0.023 | -0.015 | 0.062 | RCP45:TmaxC2 | -0.023 | -0.062 | 0.015 |
| RCP45:TmaxC3 | 0.003 | -0.027 | 0.034 | Paleo:TmaxC3 | -0.003 | -0.034 | 0.027 | Paleo:TmaxC3 | 0.000 | -0.030 | 0.030 |
| RCP85:TmaxC3 | 0.000 | -0.030 | 0.030 | RCP85:TmaxC3 | -0.003 | -0.034 | 0.027 | RCP45:TmaxC3 | 0.003 | -0.027 | 0.034 |
| RCP45:TmeanN2 | **0.058** | **0.019** | **0.098** | Paleo:TmeanN2 | **-0.058** | **-0.098** | **-0.019** | Paleo:TmeanN2 | **-0.067** | **-0.107** | **-0.027** |
| RCP85:TmeanN2 | **0.067** | **0.027** | **0.107** | RCP85:TmeanN2 | 0.009 | -0.031 | 0.049 | RCP45:TmeanN2 | -0.009 | -0.049 | 0.031 |
| RCP45:TmeanR2 | 0.002 | -0.025 | 0.029 | Paleo:TmeanR2 | -0.002 | -0.029 | 0.025 | Paleo:TmeanR2 | -0.010 | -0.037 | 0.018 |
| RCP85:TmeanR2 | 0.010 | -0.018 | 0.037 | RCP85:TmeanR2 | 0.007 | -0.020 | 0.035 | RCP45:TmeanR2 | -0.007 | -0.035 | 0.020 |
| RCP45:SoilmoistC1 | 0.001 | -0.027 | 0.028 | Paleo:SoilmoistC1 | -0.001 | -0.028 | 0.027 | Paleo:SoilmoistC1 | -0.008 | -0.035 | 0.020 |
| RCP85:SoilmoistC1 | 0.008 | -0.020 | 0.035 | RCP85:SoilmoistC1 | 0.007 | -0.021 | 0.034 | RCP45:SoilmoistC1 | -0.007 | -0.034 | 0.021 |
| RCP45:SoilmoistN1 | 0.016 | -0.013 | 0.045 | Paleo:SoilmoistN1 | -0.016 | -0.045 | 0.013 | Paleo:SoilmoistN1 | -0.002 | -0.031 | 0.027 |
| RCP85:SoilmoistN1 | 0.002 | -0.027 | 0.031 | RCP85:SoilmoistN1 | -0.014 | -0.043 | 0.014 | RCP45:SoilmoistN1 | 0.014 | -0.014 | 0.043 |
| RCP45:HabitatM | -0.073 | -0.188 | 0.041 | Paleo:HabitatM | 0.073 | -0.041 | 0.188 | Paleo:HabitatM | **0.145** | **0.030** | **0.259** |
| RCP85:HabitatM | **-0.145** | **-0.259** | **-0.030** | RCP85:HabitatM | -0.071 | -0.186 | 0.043 | RCP45:HabitatM | 0.071 | -0.043 | 0.186 |
| RCP45:HabitatML | -0.024 | -0.134 | 0.086 | Paleo:HabitatML | 0.024 | -0.086 | 0.134 | Paleo:HabitatML | 0.065 | -0.045 | 0.174 |
| RCP85:HabitatML | -0.065 | -0.174 | 0.045 | RCP85:HabitatML | -0.041 | -0.151 | 0.069 | RCP45:HabitatML | 0.041 | -0.069 | 0.151 |

**Table S7 | Range loss and range gain have low phylogenetic signal for all time periods**: paleo (6000 mya) to present (1960-1990); present to two future scenarios (2070; RCP4.5 and RCP8.5); Blomberg's K and Pagel's λ and p-values for hypothesis that K and λ are different from 0.

| | K | *p*-val | λ | *p*-val |
|---|---|---|---|---|
| *Percent loss* | | | | |
| paleo | 0.22 | 0.44 | 0.06 | 0.65 |
| RCP4.5 | 0.20 | 0.64 | 0.00 | 1.00 |
| RCP8.5 | 0.22 | 0.49 | 0.00 | 1.00 |
| *Percent gain* | | | | |
| paleo | 0.16 | 0.79 | 0.00 | 1.00 |
| RCP4.5 | 0.21 | 0.62 | 0.07 | 0.75 |
| RCP8.5 | 0.23 | 0.59 | 0.00 | 1.00 |

SUPPLEMENTARY FIGURES

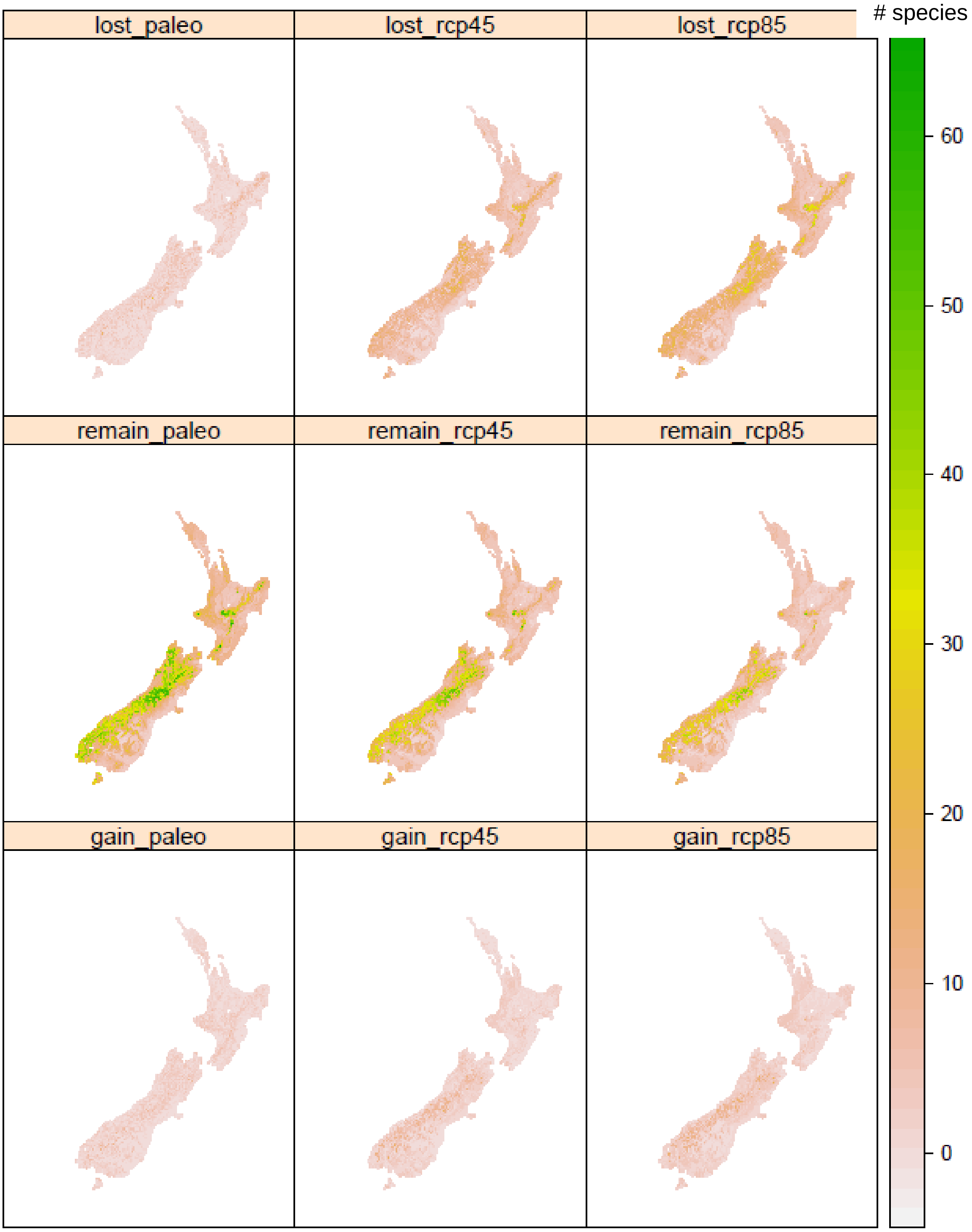


**Figure S1 | Overall patterns of range loss, retention, and gain.** Maps show number of species in each 1-km grid cell experiencing projected range loss (row 1), retention (row 2), or gain (row 3) between mid Holocene (6 kya; paleo) and present (1960-1990) and between present and future scenarios (2070; "rcp45" and "rcp85") for 93 species of *Veronica*.

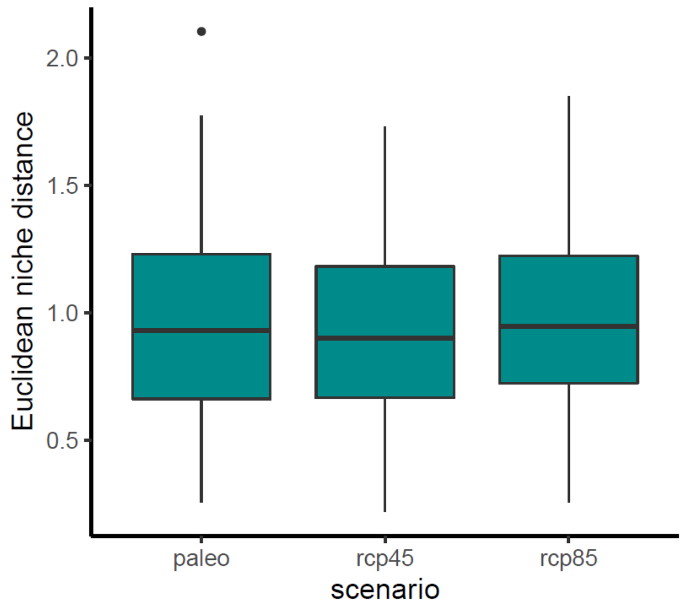


**Figure S2 | Niche change between time periods is similar in past and future.** Euclidean distance between 17 modelled physiological traits in mid-Holocene (6 kya; "paleo") and present climate (1960-1990) and between present and future scenarios (2070; "rcp45" and "rcp85") for 93 species of *Veronica*. Boxes show median and quartiles; whiskers show minimum and maximum, and black point shows an outlier >1.5-times the interquartile range.
